## Supplementary Information for "Mesoscale phase separation of chromatin in the nucleus"

#### I. MODELING THE STRETCHING ENERGY AND BENDING ENERGY OF THE CHROMATIN CHAIN

The stretching energy of the chromatin chain is calculated as:

$$E_s^{\text{chrom}} = \sum_i k_s(\mathbf{r}_i - 2a)^2 \tag{1}$$

where $\mathbf{r}_i$ is the distance between $i^{th}$ and $(i+1)^{th}$ beads. $k_s$ and $2a$ are respectively, the spring constant and equilibrium distance between two neighboring beads.

The chromosome has a persistence length of 2 beads (1.2 kb) which is based on a previous estimate [1] of a persistence length of 1-2 kb for interphase chromatin. Since the bending energy determines the persistence length, this estimate allows us to write the bending energy of chromosome as:

$$E_b^{\text{chrom}} = \sum_i k_b(1 - \cos\theta_i) \tag{2}$$

where $\theta_i$ is the relative angle between two neighboring beads, $k_b$ is bending stiffness which is taken as $l_p k_B T/\sigma$. Here $l_p$ is persistence length, $\sigma$ is diameter of the bead, $k_B$ is Boltzmann constant, and T is absolute temperature.

### II. SIMULATION PARAMETERS

| Para-meter | Description | reduced unit | SI unit |
|---|---|---|---|
| $k_BT$ | Thermal energy | 1.0 | $4.1 \times 10^{-21}$J |
| $m$ | Bead mass | 1.0 | $10^{-21}$ kg |
| $\sigma$ | LJ size parameter | 1.0 | 10 nm |
| $\epsilon$ | LJ energy parameter | $1.0k_BT$ | $4.1 \times 10^{-21}$J |
| $r_c$ | Contact distance | $2.5\sigma$ | 25 nm |
| $k_s$ | spring constant | $100k_BT/\sigma^2$ | 0.41 Jm$^{-2}$ |
| $l_p$ | persistence length | $2.0\sigma$ | 20nm |
| $k_b$ | Bending stiffness | $2.0k_BT$ | $8.2 \times 10^{-21}$J |
| $\tau$ | damping time | $1.0 \times (3\pi\eta\sigma^3/K_BT)$ | 2$\mu$s |
| $\Delta t$ | Time step | $0.01\tau$ | 20 ns |

TABLE I: Parameter values used in the simulations.

### III. UNCONFINED CHROMATIN CHAINS: SELF-AVOIDING AND SELF-ATTRACTIVE

Here we demonstrate that in the absence of confinement, the chromatin chain of our simulation model obeys the known scaling behavior of long polymer chains in both the self-avoiding (good solvent) and self-attractive (poor solvent) scenarios. Self-avoiding chains are modeled by cutting off the LJ potential when the force goes to zero. As defined in the main text, this occurs for the cutoff $r_c = 2^{1/6} = 1.1225$. Self-attracting chains are modeled by including the attractive region of the LJ potential and the cutoff is set at $r_c = 2.5$. For a persistence length of two beads and a and LJ potential strength of $\epsilon = 1k_BT$, we simulated self-avoiding chromatin chains of $N = 100, 200, 500,$ and 1000 beads and self-attracting chains for additional values of $N$ up to 100,000 as shown in the figure. To obtain the power law scaling from our simulations, we calculated the radius of gyration $R_g$ for these various values of $N$ and write.

$$R_g = \frac{l_p}{2} N_p^{\nu} \tag{3}$$

where $l_p$ is persistence length, $N_p$ is number of persistence length ($N_p = N/l_p$), and $\nu$ is the scaling exponent. For the self-avoiding case, a good fit is found with $\nu = 0.59$ and for the self-attracting case, $\nu \approx 1/3$ as expected [2] from the theory of unconfined chains.

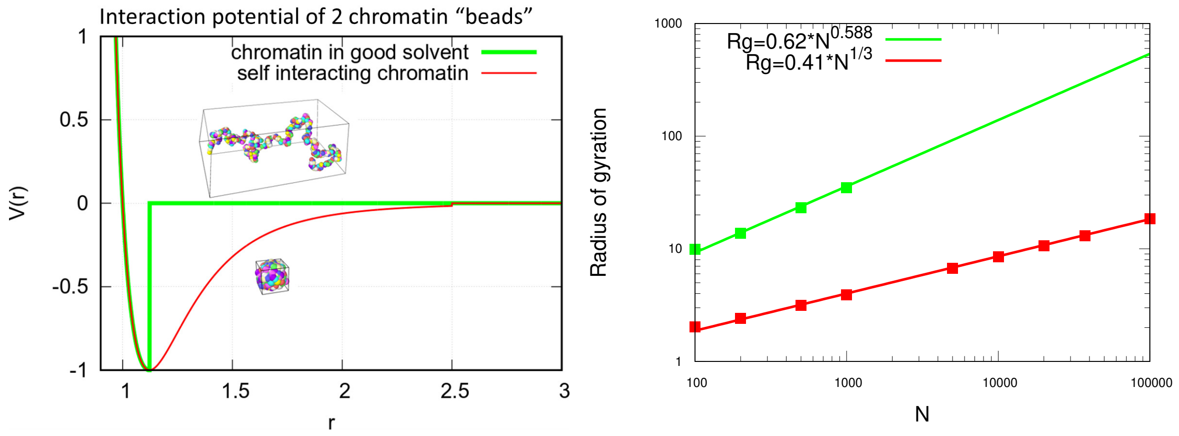


(a)

(b)

Fig. S1: (a) LJ potential as a function of the distance between two beads, with snapshot of simulations of chromatin chain having 500 beads. Snapshots show unconfined, self-avoiding chromatin in good solvent conditions for a cutoff distance $r_c = 2^{1/6} = 1.1225$ (green) and self-attractive chromatin in poor solvent conditions, for a cutoff distance $r_c = 2.5$. (b) Radius of gyration $Rg$ vs. the number of beads $N$ is calculated from the simulations of unconfined chromatin chaina having $N = 100, ...100,000$ beads (green and red dots). The green and red lines are guides to the eye. They can be fit with the power laws shown, demonstrating the polymeric scaling in the unconfined case for both good and poor solvent conditions..

### IV. ANALYSIS OF LADS SEQUENCE DATA OF *DROSOPHILA*.

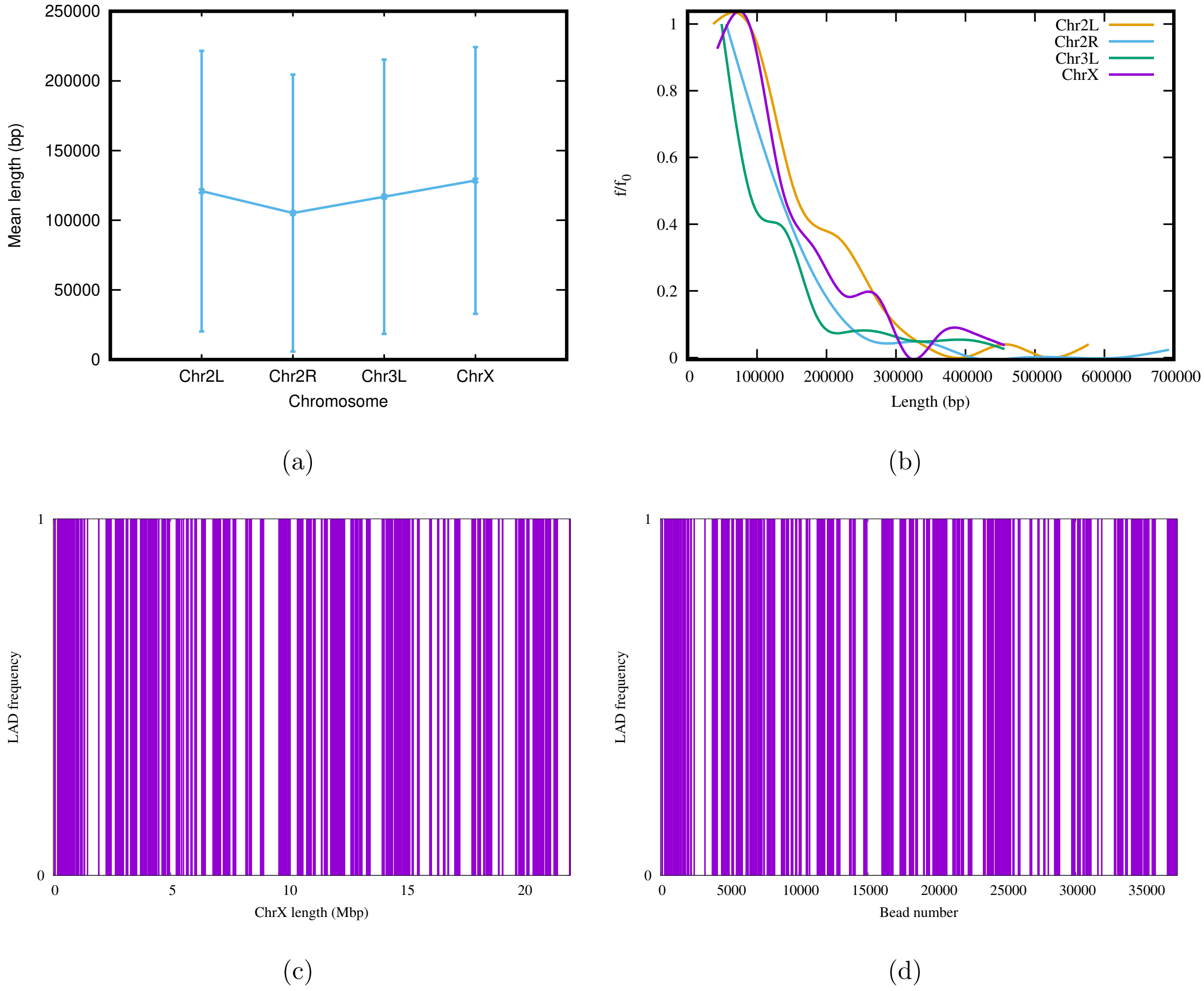


Fig. S2: (a) Mean cluster length of LAD for different chromosomes of *Drosophila* [3]. (b) Length distribution of LAD for different *Drosophila* chromosomes. In both subfigures (a) and (b), the vertical bars represent the standard deviation (SD). (c) An alternating distribution of LAD (violet) and non-LAD (white) along the 22.4 Mbp regions of chromosome X (ChrX). (d) Same LAD distribution used in our coarse-grain model with a chromatin chain of $37,333$ beads. From the sequence data it is clear, LAD regions in *Drosophila* consist of $\approx 150$ beads, and not single ones.

### V. ALTERNATE SIMULATIONS FOR RANDOM LAD SEQUENCES

To demonstrate that our simulation results do not depend on any specific LAD sequence data, we also generated LAD distributions using a Monte-Carlo method. We considered two cases for LAD binding to lamin: (i) Randomly distributed as single beads, (ii) Exponentially distributed cluster with a mean of 150 beads.

**(i) LAD randomly distributed as single beads:** In this method, we consideres a fraction $f$ of LAD beads within the chromatin chain that can bond to the lamin and distributed these randomly along the chain as single beads.

**(ii) LAD are exponentially distributed in clusters:** In this method, we used basic knowledge of the experimental sequence data for *Drosophila* that indicates that LAD are distributed along the chromatin chain in an exponential manner, in regions with a mean length of $\approx$ 150 beads. From this distribution, we used Monte-Carlo method to generate LAD regions along the chromatin chain. We first generated LAD clusters for a fraction $f$ of chromatin beads from an exponential distribution with a mean length 150 beads. We then arranged these clusters in descending order ($m1 > m2 > m3....$ where $m1, m2, m3$ are the number of beads in each cluster). Using a Monte-Carlo method, we first randomly choose a position ($1...N$ where $N$ is the total number of beads in the chain) for the cluster of size $m1$. Next, we choose a position for the cluster of $m2$ LAD beads, making sure that it does not overlap with the first cluster of $m1$ beads. After placing the $m1$ and $m2$ clusters in the chromatin chain,the $m3$ cluster can be placed between these two only if there is enough space between the two cluster to fit $m3$ beads. We repeat the process until the sum of all the clusters is equal to $Nf$ ($m1 + m2 + .... = Nf$).

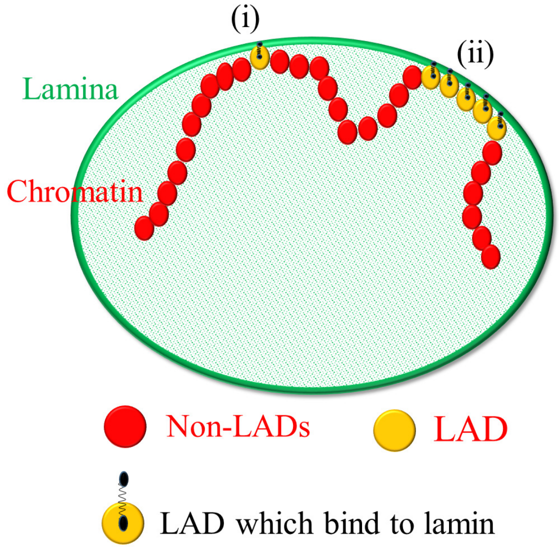


(a)

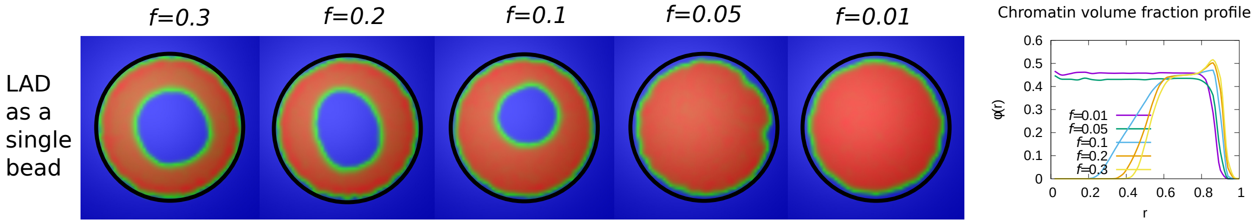


(b)

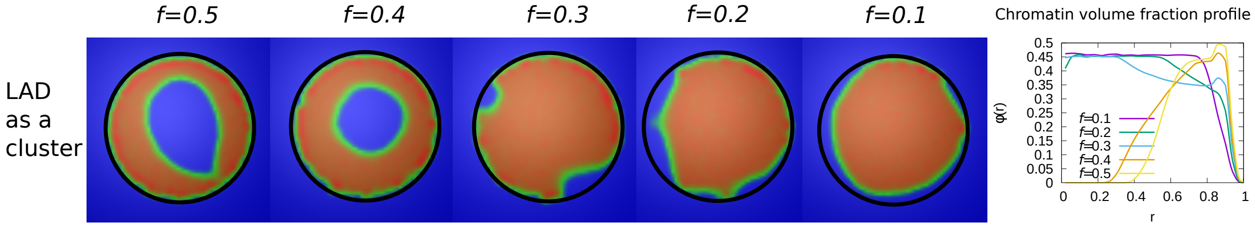


(c)

Fig. S 3: (a) Schematic diagram describing our coarse-grained model of chromatin-lamina interactions. Two cases for LAD binding to lamin: (i) Randomly distributed as single beads, (ii) Exponentially distributed cluster (many beads). Chromatin concentrations and local chromatin volume fraction profiles for $\phi = 0.3$ and $\epsilon = 1$ are shown when LAD binds to lamin as a (b) single bead (c) cluster.

### VI. LOCAL VOLUME FRACTION PROFILES OF LAD, NONLAD, AND LAMIN BEADS

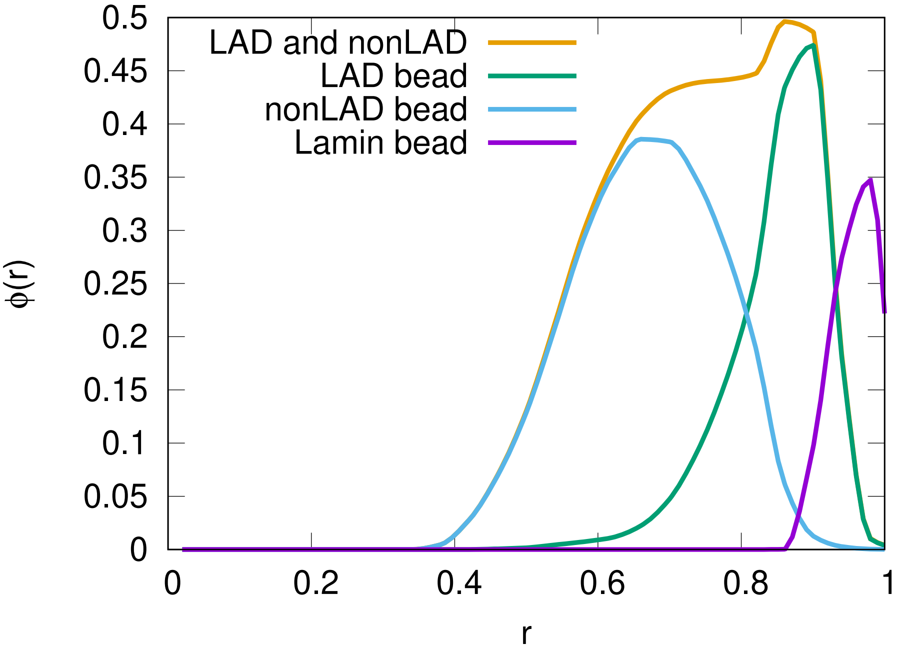


Fig. S4: For $\phi = 0.3, \epsilon = 1, f = 0.5$, the local volume fraction profiles of LAD, nonLAD and lamin beads are shown. The graph shows the location of each of these bead types in the spherical volume where $r = 0$ is the sphere center and $r = 1$ is the sphere surface.

### VII. SNAPSHOTS OF THE SIMULATED SYSTEM VISUALIZED IN TERMS OF BEADS

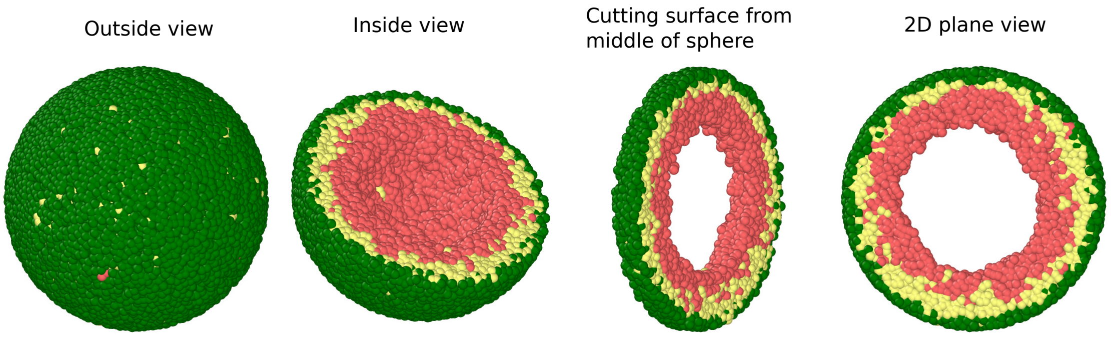


Fig. S 5: *Outside view:* The spherical volume representing the nucleus is surrounded by lamin beads (green). Within the sphere ,the chromatin chain of N=37,333 beads comprises two types beads: LAD (yellow) and non-LAD (red). *Inside view:* Spherical system is cut at the equatorial plane into two hemispheres, so that chromatin (LAD: yellow beads, non-LAD: red beads) within one hemisphere is visible. *Section cut from the equitorial plane of the hemisphere:* Sphere is cut from its equatorial plane into a slice with a width of (1/5)th of sphere diameter. *2D plane view:* We show the equatorial (xy) plane view of the 3D surface where it is clear that there is no chromatin in the middle which is filled with the aqueous phase.

### VIII. TRANSITION FROM PERIPHERAL TO CENTRAL CHROMATIN LOCALIZATION.

Here we show the transition from periphercal to central chromatin localization for small chromatin volume fraction $\phi = 0.1$ as the fraction of chromatin that can bind to the lamin is varied from $\psi = 1$ to $\psi = 0.1$.

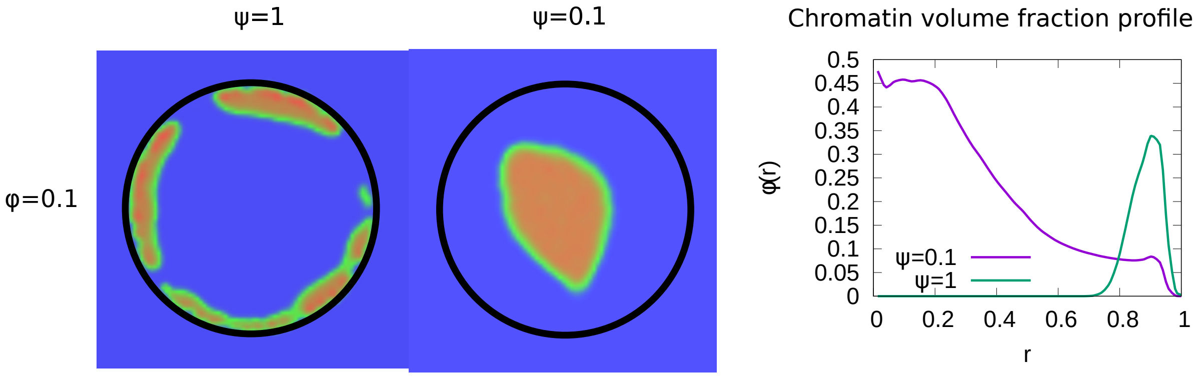


Fig. S6: *Left panel:* For a small chromatin volume fraction $\phi = 0.1$, simulation snapshots show peripheral chromatin localization for $\psi = 1$ and central localization (but see the next section on the wetting droplet) for $\psi = 0.1$. *Right panel:* Local chromatin volume fraction shows a peak near the nuclear periphery $(r = 1)$ for $\psi = 1$ while for $\psi = 0.1$, the peak of the local volume fraction is shifted towards the center $(r = 0)$.

### IX. WETTING DROPLET AND CENTRAL CHROMATIN ORGANIZATION FOR RELATIVELY WEAK LAD-LAMINA INTERACTIONS.

From simulations of the chromatin for small values of $\psi$ (fraction of LAD which can bond to the lamin), we found a central organization of chromatin with several “arms” (see “arms” in Fig.S 7(a)) that contact the lamina. This occured when the unbinding probability (related to the bond breaking as explained below) was relatively small and is thus a kinetic effect. Nevertheless, the experiments indicate such configuration. Since each LAD domain is rather large ($\approx$ 150 beads and not a single “monomer” of the chain), once it binds to the lamina, unbinding – which must involve the near-simultaneous detachment from the lamina of all 150 beads – occurs with low probability. These conditions resulted in the “arms” that resembled the experiment (see Fig.S 7 (b)). Different LAD domains will randomly contact the lamina at different positions, resulting in several “arms”. If we greatly increase probability of unbinding, then even if an entire arm randomly binds, it can unbind and the system can reach the equilibrium of an approximately wetting droplet that contacts the lamina in one region and not via many “arms”. Thus, this is a matter of time scales and the irreversibility of the LAD-lamin binding that determineds whether the chromatin in the weak-binding limit, shows central localization with several bound "arms" or whether it equilibrates to the wetting droplet. We recall that these effects refer to LAD-lamin binding as opposed to the physical chromatin-chromatin attractions, since LAD-lamin attractions are mediated by specific binding proteins. In Fig.S 7(a), we show 3D snapshots of simulations for $\psi = 0.1, \phi = 0.1, \epsilon = 1$. All the configurations show central organization of chromatin. The difference between the first two from the left is in the detachment kinetics of the LAD domains; we control this by varying the distance $r_b$ between the LAD and the lamin at which the bond between them (biophysically mediated via BAF and other proteins not explicitly included in our simulation) breaks; that is, the spring energy representing the bond, abruptly goes to zero when the bond is stretched to a distance larger than $r_b$. In the "wetting droplet" configuration (second from left) we take $r_b = 1.1225$, so that bond breaking happens much more easily than in the simulations (first from left) that show several "arms" where $r_b = 2.5$ and the LAD-lamin distance can fluctuate over a larger range and still remain bonded. Thus, for $r_b = 1.1225$, the chromatin equilibrates into a "wetting droplet" with the relatively simple geometry shown here with no "arms" that are "stuck" to the lamina. The third picture from the left shows the chromatin localization in the center with neither "arms" nor "wetting droplet" configurations for the case where no bonds are formed between the LAD and the lamin. The center of mass of the chromatin can be found anywhere within the nuclear volume.

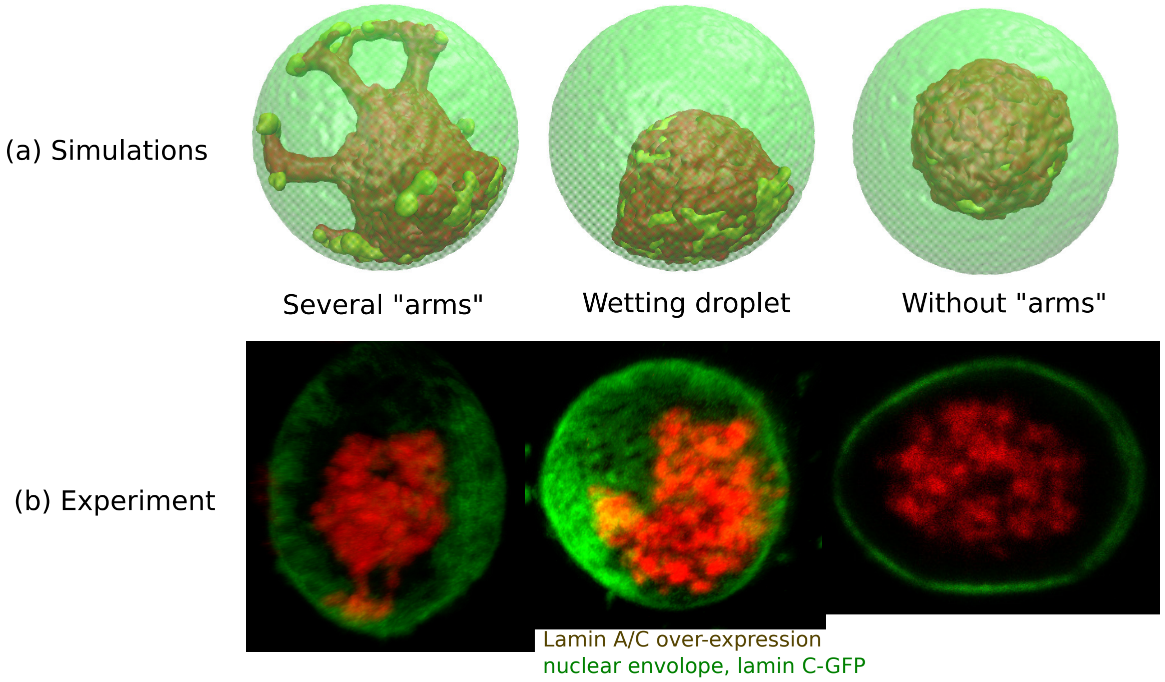


Fig. S7: (a) 3D snapshots of simulations showing central organization of chromatin with/without "arms" as well as the wetting droplet. (b) Experiments also suggest central organization of chromatin with/without "arms" as well as the wetting droplet for the case of Lamin A/C overexpression [4].

### X. COMPARISON OF THE SIMULATIONS WITH THE EXPERIMENTAL IMAGES

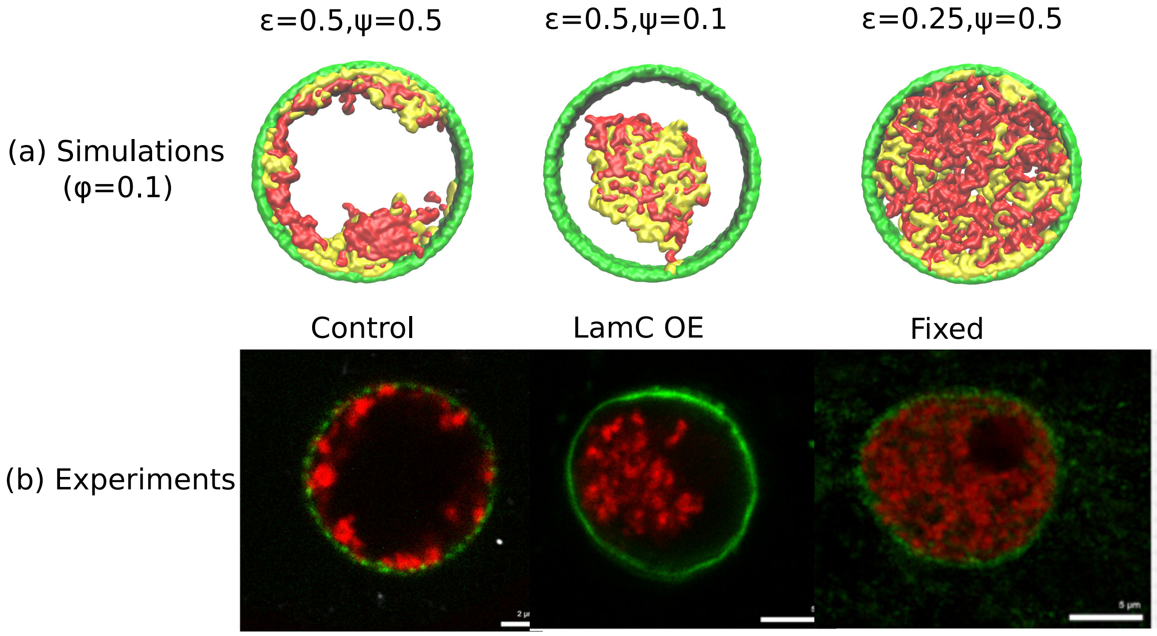


Fig. S 8: (a) Snapshots of simulations for a global volume fraction of chromatin $\phi = 0.1$. For relatively high chromatin-chromatin attractions ($\epsilon = 0.5$) and relatively high chromatin-lamina binding interactions ($\psi = 0.5$), the simulation shows peripheral organization of chromatin. For $\psi = 0.1$ (small) and $\epsilon = 0.5$ (high), we obtain central organization of chromatin (see previous section on the wetting droplet). For $\psi = 0.5$ (high) and $\epsilon = 0.25$ (small), the simulations show conventional organization of chromatin, where the entire nucleus is filled. (b) Different organization of chromatin obtained in the experiments: peripheral, central and conventional are seen in experiments in the control, lamin A/C overexpression and fixed nuclei respectively [4].

[1] J. F. Marko and E. D. Siggia, Molecular biology of the cell **8**, 2217 (1997).

[2] M. Rubinstein, R. H. Colby, et al., *Polymer physics*, vol. 23 (Oxford university press New York, 2003).

[3] J. W. K. Ho, Y. L. Jung, T. Liu, B. H. Alver, S. Lee, K. Ikegami, K.-A. Sohn, A. Minoda, M. Y. Tolstorukov, A. Appert, et al., Nature **512**, 449 (2014).

[4] D. Amiad-Pavlov, D. Lorber, G. Bajpai, S. Safran, and T. Volk, bioRxiv (2020).
